# Persistent, Multi-Species Outbreaks and Long-Range Transmissions of a Measles-Like Virus

**DOI:** 10.64898/2026.07.20.739659

**Authors:** Nicole Nova, Katherine A. Solari, Kimberlee B. Beckmen, Martin Gilbert, Ellen E. Brandell, Julia A. Palacios, Elizabeth A. Hadly, Erin A. Mordecai, Dmitri A. Petrov

## Abstract

Canine distemper—a measles-like disease with high mortality and presently without a cure—poses a major threat to wild and domestic carnivores globally. Domestic dogs are generally considered to be the main reservoir and long-range transmitter of the disease, but the role of wildlife is likely underestimated and the long-term persistence of CDV in wildlife has never been assessed. We sequenced canine distemper virus (CDV) full and partial genomes that were sampled over a decade (2012–2021) from Arctic foxes and other canids in Alaska and Yellowstone, and compiled a dataset of all published CDV genomes sampled across 32 species globally. We show the first ever evidence of persistence of CDV in wildlife (for almost a decade) with explosive transmission dynamics crossing host species barriers. Strains sampled from the Arctic for the first time connect North America to Eurasia, and are distinct from the Yellowstone strain and other known North American lineages. This suggests that separate wildlife outbreaks occur concurrently in North America, with possible introductions from Eurasia. The long-term persistence, long-range movement and explosive spread of this devastating panzootic virus that we document within wildlife is alarming and highlights the need for increased monitoring efforts to better protect wildlife populations globally.

## 1 Main text

Canine distemper virus (CDV) likely originated when measles spilled over from humans to domestic dogs in the 18th century (*1*). It is a lethal virus that has been detected and sequenced in domestic and wild mammals worldwide across four Orders—Rodentia, Primates, Artiodactyla, and most commonly Carnivora (*2*), and poses an existential threat to several endangered species (*2–4*). Theoretically, measles-like viruses that cause acute illness and lifelong immunity can only persist in a population if the influx, number, and density of susceptible hosts is large—suggesting that for CDV, domestic dogs may be a more likely reservoir than the now smaller populations of wild carnivores. Thus, previous studies posited that CDV is maintained in domestic dogs with occasional spillover events into wildlife (*5*, *6*). However, there have been recent reports of high mortality events, high seroprevalence, and evolving strains of CDV in wildlife, suggesting the existence of CDV reservoirs outside of domestic dogs (*7–10*). Additionally, most previous studies showed that the CDV phylogeny is split into continental lineages, suggesting diffuse transmission dynamics among species within continents (these phylogenies were based on a single gene [i.e., the “H” gene]) (*5*, *6*, *11*). However, more recent studies, many of which used full or partial viral genomes, show phylogenies with mixed continental origin, suggesting cases of cross-continental transmission and multiple continental introductions (*12–14*). Previous studies hypothesized that these global transmission events are caused by domestic canines transported by humans (*12*, *15*); however, the role of long-range transmission by wildlife can not be ruled out. Consequently, there is reason to believe that current CDV control efforts, which focus largely on domestic dogs, may be underestimating the role of wildlife in local and global transmission patterns and dynamics.

In order to gain a better understanding of the role of wildlife in CDV persistence and transmission, we must assess if CDV can be maintained in a wildlife population and the temporal and spatial dimensions of wildlife outbreaks. Identifying potential wildlife reservoirs requires dense sampling of wildlife in a region over an extended period of time paired with whole genome sequencing of CDV to follow viral transmission.

Here, we carried out such a study by capitalizing upon existing wildlife tissue samples from Alaska spanning almost a decade as well as one sample from Yellowstone National Park, Wyoming, USA. We generated CDV whole genome sequencing data from these samples and used these data to investigate local outbreak dynamics across Alaska. We also leveraged all existing CDV genome sequencing data to put these newly generated sequences into a global context and assess global transmission dynamics.

We extracted RNA from a total of 235 carnivore samples (Table S1). Samples consisted of brain tissue originally collected as part of rabies surveillance efforts, or collected from road kills or carcasses of unknown cause of death. These samples spanned a nine year period (2012–2021) and more than a 1,000 km range in mostly northern Alaska. We also processed one wolf (*Canis lupus*) sample from Yellowstone National Park collected in 2017. After extracting RNA, we screened samples for CDV using a PCR-based method. We then employed a novel short-read multiplex PCR approach to pursue whole genome sequencing from all CDV-positive samples. Ultimately, we sequenced 46–97% (6,961–14,652 bp) of the CDV genome (on average 72% of the genome) from 34 canids: 32 Arctic foxes (*Vulpes lagopus*) and one domestic dog from Alaska, and one wolf from Yellowstone (Table 1). First, we focused on the CDV sequence data generated from the Alaska sample set and applied Bayesian phylogenetic and phylodynamic analyses to assess the relatedness among these sequences, visualize the spread of this outbreak through time, and calculate outbreak parameters. We next compiled and analyzed a global dataset containing our newly sequenced viral sequences (from Alaska and Yellowstone) and all previously published CDV genomes (n = 249) from numerous species (n = 32)—including canids, felids, mustelids, primates, and pinnipeds. We used this global tree to assess how the CDV sequences that we generated from Alaska and Yellowstone relate to each other and to other published data, as well as to visualize the evolution and global spread of this panzootic virus through time.

**Table 1.** The number of tissue samples, CDV-positive samples, and successfully sequenced CDV genomes (binned by coverage) obtained in this study per species and sample year. Here we show a subset of species and sample years that yielded successful CDV genome sequences from a larger CDV screening effort in Alaskan carnivores and Yellowstone National Park gray wolves (Table S1). The game management units (GMUs) numbers indicate the GMU unit numbers (shown in Figure 1a) from which the samples originated (and not the number of unique GMUs). CDV stands for canine distemper virus.

| Species | Year | Total no. of samples processed | CDV detected | >70% genome sequenced | 45–70% genome sequenced | GMU number |
| --- | --- | --- | --- | --- | --- | --- |
| Arctic fox | 2012 | 17 | 9 | 4 | 3 | 26 |
| Arctic fox | 2013 | 11 | 4 | 3 | 0 | 26 |
| Arctic fox | 2014 | 21 | 11 | 3 | 2 | 26 |
| Arctic fox | 2021 | 45 | 29 | 9 | 8 | 22 |
| Domestic dog | 2021 | 2 | 2 | 0 | 1 | 22 |
| Gray wolf | 2017 | 1 | 1 | 0 | 1 | Yellowstone |
| **Total** |  | **97** | **56** | **19** | **15** |  |

Bayesian phylogenetic analyses of the Alaska CDV sequences revealed long-term persistence of CDV with annual outbreaks in Arctic foxes in Alaska (Figure 1). The dated (time-calibrated) phylogeny of the Arctic fox CDV reveals a ladder-like structure (i.e., multiple sequential clades emerging from the previous clade), suggesting that CDV has persisted in Arctic foxes for at least 9 years (2012–2021) with periodic outbreaks, where each distinct clade in the phylogeny arises from local strains the year before (Figure 1). The estimated time to the most recent common ancestor (tMRCA) for each of the first three clades formed by the 2012–2014 sequences was about 6 months before the sequences were sampled, suggesting that the onset of each annual outbreak occurred at about half a year prior. Further, the time between the root nodes of each cluster (i.e., inter-outbreak interval) was about one year, suggesting that CDV infections in Arctic foxes in Alaska follow seasonal dynamics, with annual outbreaks commencing around late fall or early winter. For the most recent clade formed by the 2021 sequences, the estimated tMRCA was about one year before the sequences were sampled (steel blue clade in Figure 1b), suggesting that outbreak periods may extend to a year, or alternatively be a continuation of a previous outbreak the year before. Moreover, MRCA of the 2021 clade descended from the ancestor to the 2014 clade, suggesting that CDV persisted in the Arctic foxes in Alaska from 2014 to 2021. Indeed, we detected CDV in a 2016 sample (Table S1; Figure S1), but we were unable to obtain sequences for 2015 or 2017–2020 from the relatively few samples we had available for these years. The ancestor to the 2016 sample appears after the 2014 cluster and before the 2021 cluster as expected (Figure S1), but due to the relatively low genomic coverage of the 2016 sample (20%), it was excluded from the main analyses. These results are the first to document the likely persistence and regular outbreaks of CDV in a wildlife community over almost a ten-year period.

**Figure 1.**
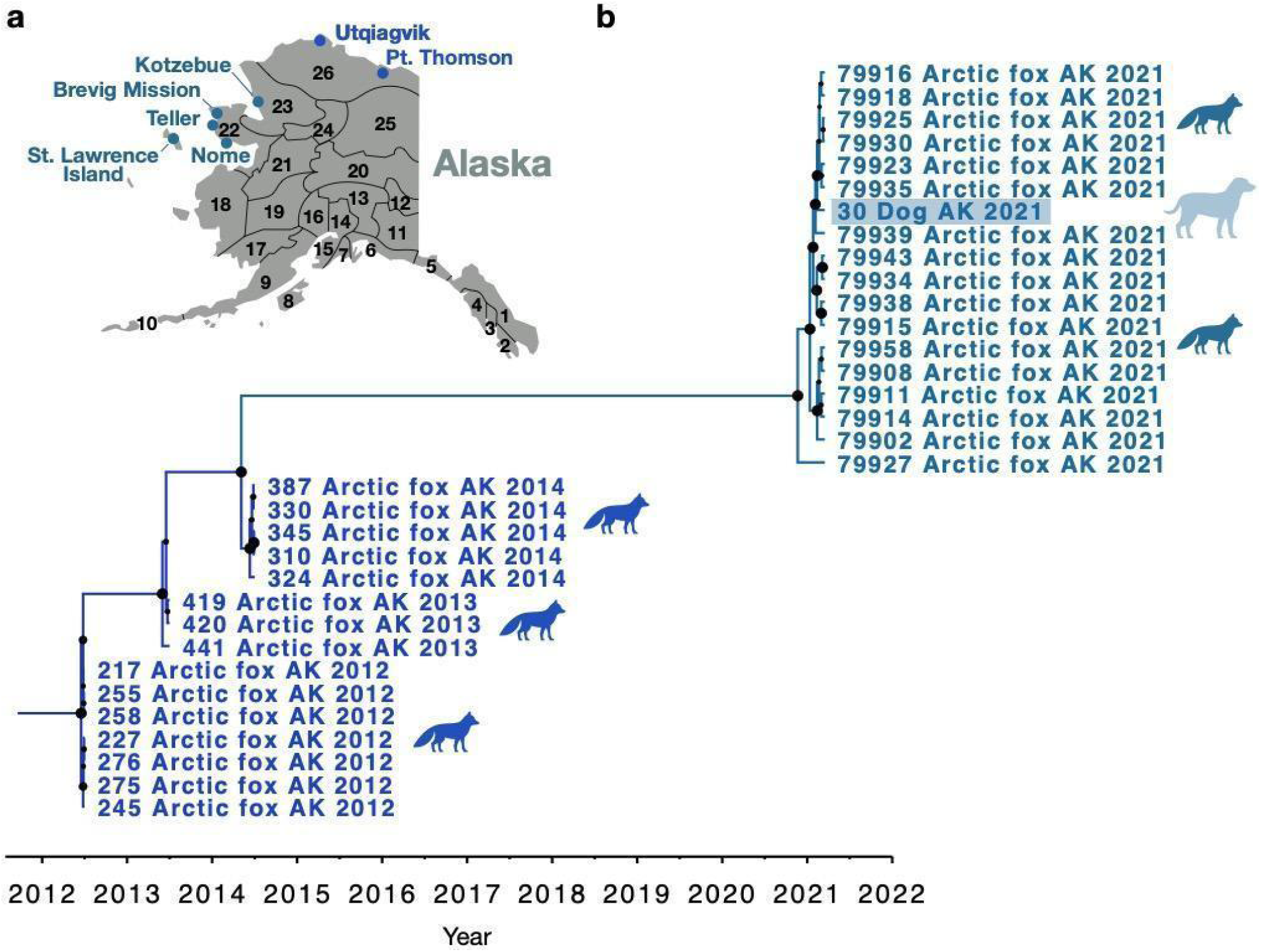
Sustained CDV transmission among Arctic foxes in Alaska for several years and cross-species transmission between Arctic fox and dog in the 2021 outbreak. (a) Map of Alaska with game management unit (GMU) locations, where most of the sampling has been done in Alaska. The colors of the samples in (b) match the colors on the map and indicate where the samples have been collected. (b) A dated phylogenetic (maximum clade credibility) tree of 33 CDV genomes from Arctic foxes and one domestic dog (highlighted in light blue) sampled in 2012–2014 (blue; bottom) and 2021 (steel blue; top) in Alaska. The horizontal positioning of each node (along the timeline) represents the estimated median of the time of divergence in years (from the most recent common ancestor [MRCA]). The size of the black dots on the nodes represents the size of the posterior probability (PP) with the maximum size (PP = 1) equivalent to the size of the black dot at the root of the tree (January, 2012). Tip labels contain animal ID for sequences obtained in this study, species common name, and sample collection year.

We also found evidence for cross-species transmission of CDV in Arctic Alaska during this single outbreak. The 2021 clade of the Arctic Alaska phylogeny (steel blue, Figure 1b) shows that the viral sequence from a domestic dog is embedded in the cluster of Arctic fox CDV genomes. This shows that domestic dogs can be part of the transmission chain of CDV in an outbreak in wildlife, not only as donors but potentially also as recipients (Figures 1b). Additionally, the sequence obtained from a red fox (*Vulpes vulpes*) in Alaska in 2021 is also embedded in the 2021 Arctic fox cluster of sequences (Figure S1). This sample only yielded ∼5% genome coverage and was thus not included in the main analyses; however, this preliminary result indicates that transmission across the different fox species could be a possibility and should be investigated further. Thus, the disease dynamics of an endemic CDV outbreak may not be governed by a population of a single species, but rather by a heterogeneous susceptible community composed of multiple species.

Bayesian phylodynamic analyses of the 2021 outbreak further revealed explosive dynamics with onset in early winter and peaking in February/March (Figure 2). The *R_0_* estimate (i.e., the average number of individuals an infected individual will infect in a fully susceptible population) for this outbreak was 7.9 on average (median: 7.5; 95% highest posterior density (HPD) [0.3, 14.9]) (Figure 2b). This is on par with the *R_0_* of the Omicron SARS-CoV-2 variant with an *R_0_* of 9.5 (*16*) and just below human measles *R_0_* of 12–18 in an unvaccinated population (*17*). This estimate, although high, is reasonable given that a previous study estimated *R_0_* for CDV to be 5.69 (± 0.67) in hyenas (*18*). The estimated effective population size *N_e_* of CDV, which is proportional to the incidence of CDV infections (outbreak size), peaked sometime between March 17 and February 27, 2021 (Figure 2a). The onset and rapid growth of *N_e_* is consistent with expectations of outbreak dynamics based on the structure of the phylogeny (steel blue clade in Figure 1b) and CDV outbreak dynamics observed in other studies (*5*, *19*, *20*).

**Figure 2.**
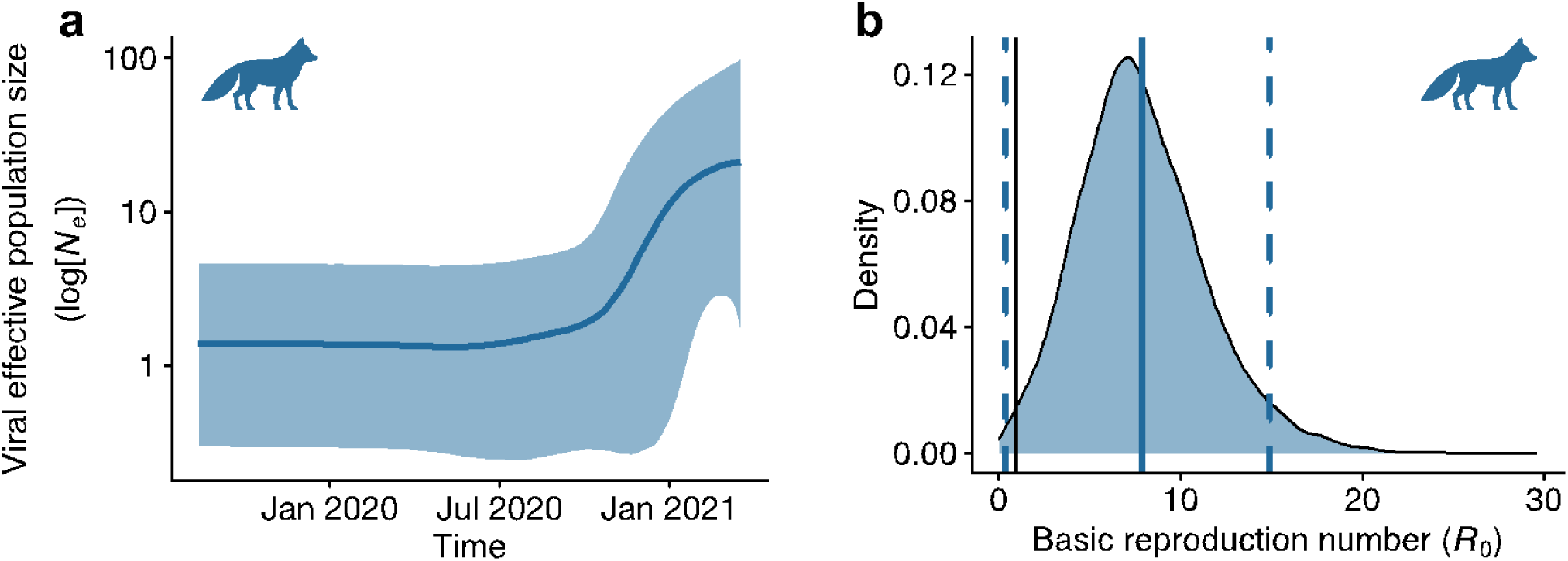
The 2021 outbreak size increased rapidly with high transmission of CDV in Arctic foxes in Alaska. (a) A Bayesian skyline plot showing the inferred effective population size, *N_e_*, of CDV, which is proportional to the outbreak size of the 2021 CDV outbreak in Alaska. The x-axis is time in years, and the y-axis is a logarithmic scale of the effective population size of CDV (proportional to the incidence). The blue solid line represents the mean, and the blue shaded region represents the 95% highest posterior density (HPD) interval. The peak mean value was 20.9 (95% HPD [1.7, 99.5]) and occurred on March 17, 2021. The peak median was 12.0 (95% HPD [2.9, 85.3]) and occurred on February 27, 2021. (b) Inferred kernel density estimate of the basic reproduction number (*R_0_*) of CDV for the 2021 outbreak in Alaska. Solid blue vertical line represents the mean (7.9; median: 7.5), the blue dashed lines represent the 95% HPD interval (0.3 and 14.9). The solid black vertical line represents *R_0_* = 1.

In order to assess how these newly sequenced Arctic strains relate to the strains from other parts of the world and to trace the global spread and evolution of this panzootic virus through time, we compiled and analyzed a global dataset containing our viral sequences and all previously published CDV genomes. The Alaska CDV genomes sequenced here are the first-ever whole genome sequences from the Arctic, and we used their positioning in the global tree to posit whether they form a cross-continental path between the Americas and Eurasia. Note that Arctic-like CDV strains were first identified in dogs in Italy (*21*), followed by dogs in the United States (*22*), and more recently in other European countries (*23–25*), and were called Arctic-like due to their H-gene sequences grouping with strains recovered from canines in Greenland in 1988 (*26*) and from a Baikal seal (*Pusa sibirica*) in Russia (*27*). However, no whole genome data for CDV from the Arctic existed until now. Our global tree shows that the Alaskan clade (blue in Figure 3) is closely related to Eurasian sequences (Figure 3) sampled from an Amur tiger (*Panthera tigris altaica*), a Baikal seal (*28*), and wolves in Italy (*29*, *30*), suggesting possible transmission between the Americas and Eurasia via the Arctic, and underscoring cross-carnivore infection dynamics.

**Figure 3.**
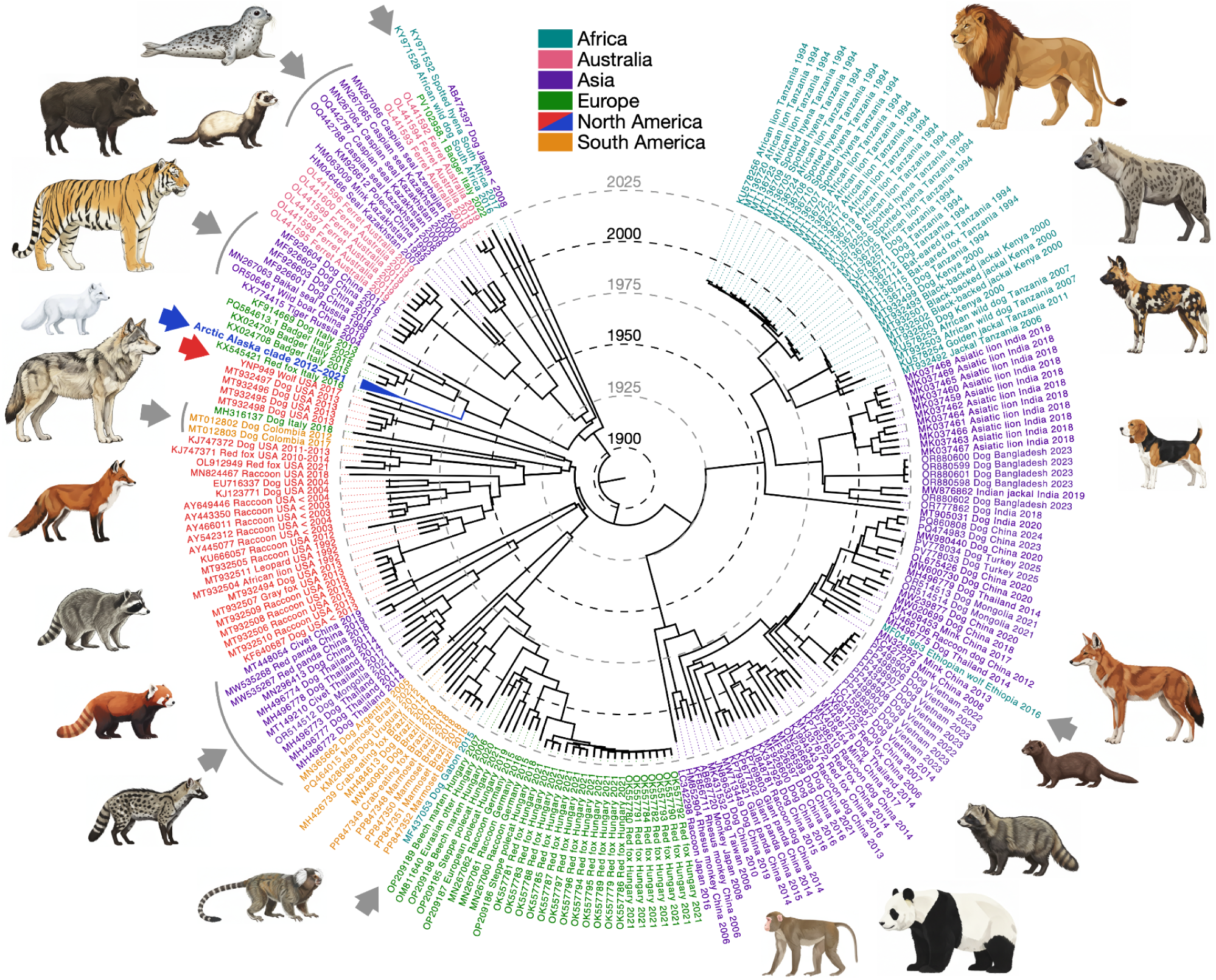
Global phylogenetic tree of existing CDV whole genome sequences. A dated phylogenetic (maximum clade credibility) tree of all previously published 215 whole genomes and the 34 genomes from this study (the Arctic Alaska clade and the Yellowstone wolf sequence are indicated by blue and red arrows, respectively). Tips are colored by continent: Africa in teal, Australia in pink, Asia in purple, Europe in green, North America in red (contiguous US) or blue (Alaska), and South America in orange. Tip labels contain NCBI Genbank accession numbers for published sequences or animal ID for sequences obtained in this study, species common name, country, and sample collection year. For published samples that did not indicate the year of sampling, we included the year they were added to NCBI, and the symbol “<” thus indicates that a sample has been collected that same year or earlier. Dashed lines in a circle represent the timeline in years. Gray arrows point to examples of cross-continental transmission. Animal icons created by NN.

This global tree also revealed that CDV clusters more by geography than by host species (Figure 3); however, CDV is not entirely split into continental lineages. Our global phylogeny suggests that multiple recent trans-continental transmission events have occurred, as some sequences or clusters are embedded in or stemming from clades of other continents (Figure 3). Indeed, the long divergence time between the Alaska clade and the Yellowstone wolf sequence (indicated by blue and red arrows, respectively, in Figure 3), as well as between other sequences from North America (red clusters in Figure 3), indicates numerous concurrent outbreaks of distinct origin in North America. This suggests the occurrence of multiple, isolated epizootics in different locations of North America, with the possibility that some, including the Alaskan clade, may have become enzootic (Figure 1b).

Our global phylogeny further allowed us to investigate the origin of all sequenced CDV genomes. Our Bayesian phylogenetic analyses placed the most probable occurrence of tMRCA at the turn of the 20th century (mean: 1896; median: 1902) with a 95% HPD interval ranging from 1844 to 1940 (Figure 3). Indeed, previous analyses using the H-gene estimated the occurrence of the MRCA to 1868 (*12*) and 1886 (*6*). Further, we estimated the evolutionary rate of the virus to be 5.9 × 10^-4^ nucleotide substitutions per site per year on average (95% HPD: [4.5 × 10^-4^, 7.3 × 10^-4^]), which is similar to previous estimates of the CDV evolutionary rate from other data (*5*, *6*). Additionally, our global phylogeny is the first to be time calibrated, highlighting that the cross-continental transmissions happened more frequently recently, especially after the mid-1900s (Figure 3). This is evidenced by the relatively short divergence times between strains from different continents (Figure 3 gray arrows). Thus, our study indicates a more rapid and recent CDV expansion at the global scale than previously postulated, especially in the 20th century, even though it likely originated in the 18th or 19th century (*1*).

Our study provides the strongest evidence to date for long-term persistence of CDV in wildlife populations with cross-species transmission and annual, explosive outbreaks. Several aspects of Arctic fox biology suggest possible explanations for their role in the persistence of CDV and the high estimated *R_0_*. First, Arctic foxes breed annually and can yield large litter sizes (18 kits on average) (*31*) enabling higher vertical transmission from parents to kits and adding numerous new susceptible individuals every year, ensuring susceptible host availability during annual outbreaks as we observed. Second, Arctic foxes are capable of trekking thousands of kilometers across sea ice (*32*, *33*), ensuring enough movement of the virus to new locations with new susceptibles. Lastly, Arctic fox behaviour of scavenging from the same carcasses in the winter combined with the ability of CDV to survive in the environment for up to two weeks in cold (∼5°C) shady places (*34*) may facilitate transmission among multiple individuals across and within species without direct contact.

Although these features of Arctic fox support their potential as a reservoir species, our results also show that this outbreak includes multiple species. This highlights how the cross-species transmission dynamics of CDV could also play a role in ensuring a large enough susceptible host community, composed of multiple species, to sustain the virus over long periods and become enzootic. A CDV outbreak involving multiple host species has been hypothesized but never definitively demonstrated in North America. For example, in the Yellowstone ecosystem, there have been several CDV outbreaks documented in wolves based on pup mortality and serological data (*35*, *36*); however, the Yellowstone wolf population alone is too small to sustain CDV, suggesting that a CDV lineage may be persisting within the multi-species community in the Yellowstone ecosystem (*37*). In the Serengeti National Park, Tanzania, Weckworth *et al*. demonstrated, using viral genome sequencing, that cross-species transmission of CDV among hyenas (*Crocuta crocuta*) and lions (*Panthera leo*) occurred during the 1992–1994 outbreak (*38*). Frequent cross-species transmission is also supported by our global tree, which shows that CDV strains cluster more by geography than host species, consistent with previous phylogenetic analyses (*5*). Thus, it seems that CDV often overcomes host species barriers in a given region and our Alaska phylogeny provides the first specific evidence of this occurring in North America.

Our global phylogenetic tree not only shows CDV moving readily among different species, but also across different continents, as previously identified by others (*12*, *39*). While previous studies hypothesize that these global transmission events are caused by domestic canines transported by humans (*12*, *15*), the data that we present show that Arctic foxes can also be responsible for long-range transmission. As mentioned above, Arctic foxes can disperse long distances, and indeed, we document the spread of a CDV strain over 1000 km presumably by Arctic foxes. Given the connectivity of continents through the Arctic by sea ice in the winter, it is feasible that Arctic foxes could be a key cross-continental transmitter of the disease.

This study is the first to document the long-term persistence of CDV in a wildlife population; however, this is also the first time that a longitudinal dataset has been used to investigate this. Further studies in other regions using CDV whole genomes from multiple species across multiple years are needed to confirm whether there are other wildlife populations—potentially red foxes (*40*), raccoons (*Procyon lotor*) (*41*), and others—where CDV has become enzootic, and whether frequent cross-species transmission of CDV within distinct communities is generalizable across systems. Indeed, more routine CDV testing of domesticates and wildlife performed by veterinarians globally followed by genome sequencing, using any of the numerous methods now available to do so, is pivotal to achieving a more complete understanding of the spread and evolution of this devastating pathogen. At the very least, we show how wildlife tissue samples collected opportunistically from carcasses or from animals culled with suspected rabies (that tested negative) can be repurposed to detect and sequence CDV. This approach yielded 34 full or partial CDV genomes, producing a 131% increase of CDV genomes from North America and a 16% increase of genomes globally, allowing for a more robust assessment of the within and between species transmission dynamics, regional maintenance and global distribution of CDV.

Answering some of these open questions and gaining a better understanding of CDV transmission patterns and dynamics is essential given how devastating it has proven to be for threatened and endangered wildlife populations. In recent decades, CDV outbreaks have been associated with temporary declines of 30% in Serengeti lions (*19*), 45% in Yellowstone wolves (*35*, *42*, *43*), 68% in Ethiopian wolves (*Canis simensis*) (*44*), and 95% in Channel Island foxes (*Urocyon littoralis*) (*45*), while almost completely extirpating the black-footed ferret (*Mustela nigripes*) population in Wyoming (*46*). In 2013, CDV was detected in the endangered Amur tigers and Amur leopards (*Panthera pardus orientalis*), with only 500 and 60 individuals left in the wild, respectively (*47*). Understanding the dynamics of how CDV spreads between continents and between species is critical to support conservation of vulnerable populations. Vaccines could be a promising tool in protecting endangered wildlife populations from CDV (*48*). However, while numerous types of vaccines currently exist, they target a historic CDV strain that is now 10% different from current circulating strains, underscoring the rapid evolution of CDV and suggesting that nimble vaccine design is essential (*13*). This discrepancy could explain examples of vaccine inefficacy in dogs (*49*). Questionable vaccine efficacy in domestic animals is overshadowed by the uncertain safety or efficacy in wildlife where very little testing has been done. An updated vaccine targeting currently circulating strains with confirmed safety and efficacy that can be realistically administered to wildlife populations (i.e., oral vaccines) has already been identified as an important goal moving forward (*50–53*). Should such a vaccine be developed, a better understanding of CDV wildlife reservoirs and outbreak dynamics, through screening efforts similar to what we present here, will also be pivotal to optimize when and where vaccines should be administered for the most impact.

Although the existence of wildlife reservoirs of CDV has been previously suggested (*7–10*), the longitudinal data to test this hypothesis have heretofore been unavailable. The results we present here document, for the first time, long-term persistence of CDV in wildlife with recurring explosive, multi-species outbreaks regionally. Our global analyses indicate that long-range transmission events occur frequently enough that concurrent outbreaks on the same continent, as in our study, or even the same country (*39*) may result from independent introductions rather than local spread. The combination of frequent cross-continental transmission of CDV with its ability to cause fatal, explosive, cross-species outbreaks locally is alarming and highlights the imminent threat that CDV poses to wild and domestic carnivore populations globally. Increased surveillance with CDV genome sequencing is essential to better understand the spatiotemporal dynamics and evolution of CDV such that measures can be taken to protect at-risk populations.

## 2 Methods

### 2.1 Collection of wildlife samples

Tissue samples from wild carnivores were collected opportunistically between 2012 and 2021 in Alaska and Yellowstone National Park, Wyoming, USA. Alaska has had multiple outbreaks of canine distemper in domestic dogs in the last two decades, and although CDV persistence in wildlife has been suspected, no published study to date has been able to confirm this (*54*). The wolf population in Yellowstone has been monitored by the U.S. National Parks Service since the wolves were reintroduced in 1995 and 1996, and four CDV outbreaks have been documented in the Yellowstone wolf population (1999, 2005, 2008, and 2017) based on pup mortality and serological data (*35*).

Brainstems from carnivores in Alaska (Arctic foxes, red foxes, wolves, coyotes, wolverines, black bears, brown bears, lynx, marten, and river otters) were collected between 2012 and 2021 from rabies surveillance and predator control efforts in multiple game management units (GMUs) by the Alaska Department of Fish and Game (Table S1). CDV has been detected in these carnivore species in other parts of the world (*3*, *9*, *37*, *55*). A brainstem sample from a Yellowstone wolf was obtained opportunistically from an individual that died in August 2017, for which a postmortem antigen test suggested CDV as the cause of death. All wildlife samples were transported using dry ice and stored at −80°C.

### 2.2 RNA extraction and viral sequence enrichment

To sequence CDV genomes, we first extracted RNA from the brainstem samples, most of which had unknown infection status with CDV. We then used two different methods, based on sample age and CDV RNA extraction success, as described below, to detect, target, and amplify CDV sequences. The first method was targeted enrichment using hybridization capture with myBaits, which required pre-screening of samples for CDV RNA using reverse transcription polymerase chain reaction (RT-PCR). Despite being more costly, time-consuming, and untested for CDV, the first method has proven to be effective for yielding high coverage viral genomes (e.g., for SARS-CoV-2 by (*56*)) even for samples with highly fragmented viral RNA. We thus tested this method on the older samples (Alaskan Arctic fox from 2012–2014), as RNA degradation correlates with time. We processed 8 CDV positive samples using the myBaits target sequence enrichment protocol (described in detail below). We sequenced these samples on an Illumina NovaSeq machine using paired end 150bp sequencing at Admera Health (South Plainfield, NJ). Over 3.9 million Illumina reads were generated for each sample; however, in every sample over 99.6% of these reads mapped to the Arctic fox reference genome, not the CDV reference genome (Table S2). Even though the vast majority of reads were not CDV, in one sample (ID 275) over 4,000 reads did map the CDV reference genome resulting in ∼36x coverage of ∼94% of the reference genome. We thus generated a consensus genome sequence for sample ID 275 (NCBI accession OR241432) using Analysis of Next Generation Sequencing Data (ANGSD) (*57*). Gaps in this genome were filled using a sequence from a published CDV genome that groups with Arctic CDV strains based on H-gene sequence similarity (NCBI accession KF914669) and used as a reference genome for later genome alignments. The second method was targeted enrichment using multiplex PCR, and it proved to be more effective for obtaining CDV genomes from our wildlife samples (as described below).

#### 2.2.1 RNA extraction

RNA extractions were conducted in an RNA-only PCR hood that was treated with UV light and RNAase before each use. Brainstem samples were kept frozen until sample processing. Less than 30 mg of brain tissue was cut from the sample using a razor blade and directly added to an RNAase free 2 mL tube with one 3 mm glass bead inside. Razor blades and beads were treated with RNAase before use. Samples were then processed using the Qiagen AllPrep DNA/RNA mini kit following the manufacturer’s protocol with samples kept on ice until undergoing tissue lysis for two minutes in a Biospec Minibeadbeater-96. Nulls were included in every set of extractions and resultant RNA was quantified using a Qubit RNA high sensitivity kit.

#### 2.2.2 Canine distemper virus screening and targeted enrichment using myBaits

RNA extractions from the 2012–2014 Alaskan Arctic fox samples were screened for CDV using primers from (*58*). First, single stranded complementary DNA (cDNA) was generated from RNA using the Qiagen QuantiTect Reverse Transcription Kit following the manufacturer’s protocol. Screening PCRs consisted of 12.5 μL OneTaq Master Mix (New England BioLabs), 9 μL nuclease-free water, 0.5 μL Forward Primer (10 μM), and 0.5 μL Reverse Primer (10 μM). Cycling conditions were: 94°C for 30 sec, followed by 45 cycles of 94°C for 15 sec, 54°C for 1 min, 68°C for 30 sec, followed by a final extension at 68°C for 5 min. PCR product was visualized by running 2–5 uL on a 1% agarose gel at 100 volts (gel electrophoresis) for 30 minutes. All samples used in target sequence enrichments were screened using this method; however, not all samples used in multiplex PCRs were pre-screened.

Double stranded complementary DNA (cDNA) was generated from RNA using the Thermo Scientific Maxima H Minus Double-Stranded cDNA Synthesis Kit following the manufacturer’s protocol. Resultant double stranded DNA (dsDNA) was quantified using the Qubit dsDNA high sensitivity kit and used to create Illumina libraries with the Illumina Flex library prep kit. A modified protocol using ⅕ reaction size (where all the volumes in the original protocol were divided by 5 to process 5 times as many samples per kit) was used for Illumina library preparation. Illumina libraries were then used as input for myBaits in-solution NGS library target enrichment system. Custom myBaits oligos were used to enrich the CDV sequence. Custom oligos were designed using a published CDV genome from the Arctic (NCBI accession number KF914669) as well as 14 other published CDV genomes that capture the breadth of the currently documented CDV diversity (NCBI accession numbers MH496773.1, KU578253.1, MH496779.1, AF164967.1, AB687720.2, AY386316.1, AB476401.1, KJ994343.1, AY466011.2, KF640687.1, KU578256.1, KY971528.1, KY971530.1, KU578254.1). We performed the myBaits enrichment procedure using one round of capture (24 hours at 63°C) and 16 post-capture amplification cycles.

#### 2.2.3 Canine distemper virus targeted enrichment using multiplex PCR

Primers to amplify the entire length of the CDV genome in 93 segments (Amplicon 1–Amplicon 93) using two multiplex reactions were designed using Primal Scheme online primer design tool (*59*) with a target amplicon size of 250 bp (Figure S2). The CDV genome used for primer design was the genome from sample ID 275, generated using the target sequence enrichment approach, with gaps filled in with sequence from the Arctic reference genome (NCBI accession number KF914669.1). We added adapters to these primer sequences, following the tailed amplicon method outlined in (*60*). Primers were ordered dry from Integrated DNA Technologies (IDT) and reconstituted using IDTE buffer (1xTE) pH 8. Primers were first pooled into two pools as designed by Primal Scheme (*59*). All the primers amplifying odd numbered amplicons were pooled to make one pool, the “Odd” pool, and all of the primers amplifying even numbered amplicons were pooled to make the second pool, the “Even” pool (Supplementary Files S1 and S2).

Single stranded cDNA was generated from RNA using the Qiagen QuantiTect Reverse Transcription Kit following the manufacturer’s protocol and used as input for multiplex PCR reactions. PCR protocols followed those described in (*60*). We first processed 34 unique samples (18 of which were processed in duplicate and 5 of which were processed in triplicate) and 11 nulls using multiplex PCR with two primer pools (Even and Odd). The samples were then sequenced on the MiSeq v2 platform using paired end 250 bp sequencing at Admera Health (South Plainfield, NJ). We calculated the average number of reads generated for each of the 93 amplicons using only samples for which at least one amplicon was represented by 10 reads or more (Figure S3). This included 32 samples, including 1 null for which one amplicon did produce reads. We used this data to repool primers. We split each primer pool in half by splitting up the primers that amplified well from those that did not in each pool, as described in (*59*). The half of the Even primers that worked on the most samples were pooled to make pool “Even 1” and the half of the primers that worked on the fewest samples were pooled into their own pool, “Even 2”. The Odd primers were divided into two pools, “Odd 1” and “Odd 2”, in the same way. This resulted in 4 primer pools (Even 1, Even 2, Odd 1, Odd 2). All subsequent samples were run with 4 primer pools using the same PCR methods described above.

We then processed 177 samples (23 of which were part of the set of 34 samples processed using 2 primer pools) and 26 nulls using multiplex PCR with 4 primer pools. We calculated the average number of reads generated for each of the 93 amplicons using only samples for which at least one amplicon was represented by 10 reads or more (Figure S4). This included 98 samples, including 6 nulls for which at most four amplicons were represented with 10 or more reads. Only samples with 50% or more of the genome sequenced (representing amplification of over 40 amplicons) were used in subsequent analyses. The comparison of the average number of reads generated per amplicon using 2 primer pools versus 4 primer pools is shown in Figure S5. This comparison shows that splitting the primers into 4 primer pools did improve the amplification of some amplicons; however, there are numerous regions that did not amplify well with 2 or 4 primer pools.

### 2.3 Viral sequencing and data processing

Samples prepared using the multiplex PCR method were pooled and sequenced on the Illumina MiSeq v2 platform using paired end 250 bp sequencing at Admera Health (South Plainfield, NJ). Illumina data was mapped to the reference genome (sample ID 275 filled in with a sequence from NCBI accession KF914669) using ‘bwa mem’ (*61*) and sorted using ‘samtools’ (*62*). We used iVar (*63*) to trim off primer sequences and remove any sequences less than 50 bp in length. Trimmed sequences were then sorted and indexed using ‘samtools’ (*62*). Samtools mpileup (with the –A flag to include reads that were not properly paired in variant calling) and iVar consensus (with –t 0.7 flag to only call a consensus when the dominant call made up 70% of the reads or more) were then used to call variants and generate consensus sequences. Finally, we used ‘bedtools multicoverage’ (*64*) to calculate the number of reads generated per amplicon. All sequences obtained in this study have been deposited on NCBI GenBank under accession numbers OR233558–OR233590.

### 2.4 Phylogenetic and phylodynamic analyses

The CDV genomic dataset used for phylogenetic and phylodynamic analyses consisted of the 34 CDV genomes obtained in this study, and published CDV genome sequences from the NCBI nucleotide database (http://www.ncbi.nlm.nih.gov; accession numbers shown in Figure 3). The search terms used in the NCBI database were “canine distemper virus” while specifying the sequence length to be 10,000–16,000 bp (“Canine morbillivirus”[Organism] OR canine distemper virus[All Fields]) AND (“10000”[SLEN]: “16000”[SLEN]). The search yielded 240 hits and we only included sequences with the sampled host species, geography, and year reported (either in NCBI or in corresponding publications). We also removed any fabricated vaccine strains, and sequences from experimentally infected individuals in laboratories, yielding 215 sequences (excluding those generated in our study).

Genomic sequences of CDV were aligned for strain comparisons using the Clustal Omega algorithm in the software Geneious Prime (version 2022.0.1), and later inspected manually. Appropriate nucleotide substitution models were identified for the alignment using jModeltest version 2.1.10 (*65*) using best fit model selection based on lowest Akaike information criterion (AIC) and Bayesian Information Criterion (BIC) scores. The most suitable nucleotide substitution model was the General Time Reversible [GTR] + G model (*66*).

First, we analyzed the alignment in the software TempEst version 1.5.3 to ensure that sequences were evolving in a clock-like manner (*67*). Next, the time-calibrated phylogenetic (maximum clade credibility) trees, the viral evolutionary rate, and the time to most recent common ancestor (tMRCA) were jointly inferred by fitting a molecular clock model to the sequence alignment using the software packages Bayesian Evolutionary Analysis Utility (BEAUti) and Bayesian Evolutionary Analysis Sampling Trees (BEAST) version 1.10.4 (*68*). We used the Bayesian Skyline tree model (piece-wise constant with 10 groups) with a coalescent tree prior (*69*, *70*) and an uncorrelated relaxed molecular clock model with branch rates drawn from a lognormal relaxed clock prior distribution to allow for variation in the viral evolutionary rate in each branch (*71*). Here, we used a normally distributed clock prior with mean of 4 × 10^-4^ nucleotide substitutions per site per year (s/s/y) based on the CDV evolutionary rate estimate (albeit limited to the H gene) by (*6*), with a standard deviation of 0.5 nucleotide substitutions per site per year. Nodal support of the maximum clade credibility phylogeny was estimated by calculating the posterior probability (PP).

A subset of the alignment consisting of just the Alaska strains from 2021 was used to study the epidemiology of an epizootic of CDV in wildlife (with viral sequences from 18 canid individuals). This subset of the viral genomic dataset was used to infer the viral effective population size *N_e_*(a proxy for outbreak trend) and the viral population growth rate to estimate *R_0_* of this outbreak using phylodynamic approaches described below. The idea behind phylodynamics is that viral phylogenies are shaped by both epidemiological and evolutionary processes occurring on similar time scales (*72*). Because of rapid viral mutation rates, viral phylogenies provide information about outbreak dynamics (*73*) in addition to the evolutionary history of a virus (i.e., tMRCA and evolutionary rate).

Previous studies have shown that the effective viral population size is proportional to infection incidence (*74*). For example, viral effective population size resembled the incidence of rabies in livestock (*75*). Thus, it may be used as an inferred proxy of outbreak dynamics in cases where case data is absent. The viral effective population size was inferred by fitting a Bayesian Skyline tree model (piece-wise constant with 20 groups) with a coalescent tree prior (*69*, *70*) to the 2021 sequence alignment using BEAUti and BEAST version 1.10.4 (*68*). This was performed under a strict molecular clock model and a fixed nucleotide substitution rate at the value estimated in the previous MCMC analyses (5.9 × 10^-4^ s/s/y).

The viral population growth rate (*r*) was estimated using an exponential growth tree model with a coalescent tree prior (*76*) to the 2021 sequence alignment using BEAUti and BEAST version 1.10.4 (*68*) under a strict molecular clock model and a fixed nucleotide substitution rate at the value estimated in the previous MCMC analyses (5.9 × 10^-4^ s/s/y). The growth rate parameter *r* (denoted by exponential.growthRate in the model) provides an estimate of the epidemic (or epizootic) exponential growth:

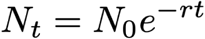

where *N_t_*is the population size at time *t* and *N_0_* is the initial population size, as *R_0_* has shown to be related to the exponential growth rate, *r*, via the Lotka-Euler equation:

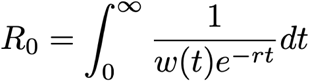

where *w(t)* is the distribution of the infection generation or incubation time (time between successive infections in a transmission chain) (*77*). Thus, *R_0_* can be inferred given estimates of *r* and the distribution of the generation time (*78*). We assumed a generation time distribution that follows a gamma distribution, assuming that the CDV generation time distribution is similar to that of other seasonal and pandemic viruses (*77*, *79*, *80*). Then,

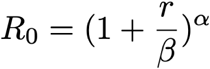

where *α* and *β* are parameters of the gamma distribution, 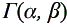, and 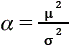 and 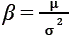, where *μ*and *σ*^2^ denote the mean and variance of the generation time, respectively. Thus, the inferred growth rate parameter distribution was transformed to an *R_0_* distribution, assuming a generation time with *μ* = 6.5 days and *σ* = 7 days in accordance with previous empirical estimates for CDV (*81–83*).

We ran four separate chains for each MCMC run. All MCMC chains were run for 10^8^ iterations (the first 10% steps were removed as burn-in). Convergence of an MCMC chain on the posterior distribution was ensured by effective sample size (ESS) values greater than 200 for each sampled parameter using TRACER version 1.7.2 (*84*). Statistical uncertainty for each parameter estimate was reflected by the mean or median value and the 95% highest posterior density (HPD).

The phylogenetic maximum clade credibility trees were summarized from the posterior distribution of trees with TreeAnnotator version 1.10.4 (*68*) and visualized using FigTree version 1.4.4 (http://tree.bio.ed.ac.uk/software/). The viral effective population size over time was summarized and visualized using the Bayesian Skyline reconstruction function in TRACER version 1.7.2 (*84*). The distributions of the inferred viral evolutionary rate and *R_0_* were visualized using the kernel density estimate function in the R package ‘ggplot2’ (*85*).

## 3 Acknowledgments

We thank Marcus W. Feldman, Giulio A. De Leo, and Paul C. Cross for helpful discussions regarding the project. We are grateful to Douglas W. Smith, Daniel R. Stahler, and Erin E. Stahler for their assistance in obtaining the Yellowstone wolf sample and the biologists from the Alaska Department of Fish and Game and the United States Department of Agriculture Wildlife Services along with Rachel Lee of the Norton Sound Health Corporation who were instrumental in obtaining the Alaska samples. We also thank Edward Dubovi for sending additional Alaska samples used previously for another project. This project was funded by the Environmental Venture Project grant from the Stanford Woods Institute for the Environment. In addition, N.N. was supported by the Philanthropic Educational Organization (PEO) Scholar Award from the International Chapter of the PEO Sisterhood, the Stanford Data Science Scholars program, the Predoctoral Fellowship from the Stanford Center for Computational, Evolutionary and Human Genomics, and the National Institutes of Health (NIH; R35GM133439). K.A.S was supported by the National Science Foundation (NSF: DEB-2225088). E.A.M. was supported by the NSF (DEB-2011147, with the Fogarty International Center), the NIH (R35GM133439), the Terman Award, the Stanford King Center for Global Development, the Stanford Woods Institute for the Environment, and the Stanford Center for Innovation in Global Health. D.A.P. was supported by the NIH (5R35GM118165-05) and Chan Zuckerberg BioHub.

## Notes

### Competing Interest Statement

The authors have declared no competing interest.

## References

1. E. W. Uhl, C. Kelderhouse, J. Buikstra, J. P. Blick, B. Bolon, R. J. Hogan, New world origin of canine distemper: Interdisciplinary insights. Int. J. Paleopathol. 24, 266–278 (2019).

2. M. Martinez-Gutierrez, J. Ruiz-Saenz, Diversity of susceptible hosts in canine distemper virus infection: a systematic review and data synthesis. BMC Vet. Res. 12, 78 (2016).

3. A. Beineke, W. Baumgärtner, P. Wohlsein, Cross-species transmission of canine distemper virus—an update. One Health 1, 49–59 (2015).

4. S. Rendon-Marin, R. Da Fontoura Budaszewski, C. W. Canal, J. Ruiz-Saenz, Tropism and molecular pathogenesis of canine distemper virus. Virol. J. 16, 1–15 (2019).

5. G. M. Ke, C. H. Ho, M. J. Chiang, B. Sanno-Duanda, C. S. Chung, M. Y. Lin, Y. Y. Shi, M. H. Yang, Y. C. Tyan, P. C. Liao, P. Y. Chu, Phylodynamic analysis of the canine distemper virus hemagglutinin gene. BMC Vet. Res. 11, 1–15 (2015).

6. Y. Panzera, N. Sarute, G. Iraola, M. Hernández, R. Pérez, Molecular phylogeography of canine distemper virus: Geographic origin and global spreading. Mol. Phylogenet. Evol. 92, 147–154 (2015).

7. R. Trebbien, M. Chriel, T. Struve, C. K. Hjulsager, G. Larsen, L. E. Larsen, Wildlife reservoirs of canine distemper virus resulted in a major outbreak in Danish farmed mink (Neovison vison). PLoS ONE 9, e85598 (2014).

8. S. Kapil, T. J. Yeary, Canine Distemper Spillover in Domestic Dogs from Urban Wildlife. Vet. Clin. N. Am. – Small Anim. Pract. 41, 1069–1086 (2011).

9. M. Tryland, A. Balboni, S. T. Killengreen, T. Mørk, O. Nielsen, N. G. Yoccoz, R. A. Ims, E. Fuglei, A screening for canine distemper virus, canine adenovirus and carnivore protoparvoviruses in Arctic foxes (*Vulpes lagopus*) and red foxes (*Vulpes vulpes*) from Arctic and sub-Arctic regions of Norway. Polar Res. 37, 1498678 (2018).

10. M. Tryland, E. Neuvonen, A. Huovilainen, H. Tapiovaara, A. Osterhaus, Ø. Wiig, A. E. Derocher, SEROLOGIC SURVEY FOR SELECTED VIRUS INFECTIONS IN POLAR BEARS AT SVALBARD. J. Wildl. Dis. 41, 310–316 (2005).

11. C. Yuan, W. Liu, Y. Wang, J. Hou, L. Zhang, G. Wang, Homologous recombination is a force in the evolution of canine distemper virus. PLoS ONE 12, e0175416 (2017).

12. H. Wang, H. Guo, V. G. Hein, Y. Xu, S. Yu, X. Wang, The evolutionary dynamics history of canine distemper virus through analysis of the hemagglutinin gene during 1930–2020. Eur. J. Wildl. Res. 69, 1–12 (2023).

13. J. Duque-Valencia, F. J. Diaz, J. Ruiz-Saenz, Phylogenomic Analysis of Two Co-Circulating Canine Distemper Virus Lineages in Colombia. Pathog. 2020 Vol 9 Page 26 9, 26 (2019).

14. Z. Lanszki, G. E. Tóth, É. Schütz, S. Zeghbib, M. Rusvai, F. Jakab, G. Kemenesi, Complete genomic sequencing of canine distemper virus with nanopore technology during an epizootic event. Sci. Rep. 12, 1–7 (2022).

15. B. Willi, A. M. Spiri, M. L. Meli, F. Grimm, L. Beatrice, B. Riond, T. Bley, R. Jordi, M. Dennler, R. Hofmann-Lehmann, Clinical and molecular investigation of a canine distemper outbreak and vector-borne infections in a group of rescue dogs imported from Hungary to Switzerland. BMC Vet. Res. 11, 154 (2015).

16. Y. Liu, J. Rocklöv, The effective reproductive number of the Omicron variant of SARS-CoV-2 is several times relative to Delta. J. Travel Med. 29, taac037 (2022).

17. R. M. Anderson, R. M. May, Infectious Diseases of Humans: Dynamics and Control (Oxford University Press, Oxford, New York, 1992).

18. S. Benhaiem, L. Marescot, M. L. East, S. Kramer-Schadt, O. Gimenez, J. D. Lebreton, H. Hofer, Slow recovery from a disease epidemic in the spotted hyena, a keystone social carnivore. *Commun*. Biol. 1, 1–12 (2018).

19. M. E. Craft, E. Volz, C. Packer, L. A. Meyers, Distinguishing epidemic waves from disease spillover in a wildlife population. Proc. R. Soc. B Biol. Sci. 276, 1777–1785 (2009).

20. M. Viana, S. Cleaveland, J. Matthiopoulos, J. Halliday, C. Packer, M. E. Craft, K. Hampson, A. Czupryna, A. P. Dobson, E. J. Dubovi, E. Ernest, R. Fyumagwa, R. Hoare, J. G. C. Hopcraft, D. L. Horton, M. T. Kaare, T. Kanellos, F. Lankester, C. Mentzel, T. Mlengeya, I. Mzimbiri, E. Takahashi, B. Willett, D. T. Haydon, T. Lembo, Dynamics of a morbillivirus at the domestic–wildlife interface: Canine distemper virus in domestic dogs and lions. Proc. Natl. Acad. Sci. 112, 1464–1469 (2015).

21. V. Martella, F. Cirone, G. Elia, E. Lorusso, N. Decaro, M. Campolo, C. Desario, M. S. Lucente, A. L. Bellacicco, M. Blixenkrone-Møller, L. E. Carmichael, C. Buonavoglia, Heterogeneity within the hemagglutinin genes of canine distemper virus (CDV) strains detected in Italy. Vet. Microbiol. 116, 301–309 (2006).

22. I. D. R. Pardo, G. C. Johnson, S. B. Kleiboeker, Phylogenetic Characterization of Canine Distemper Viruses Detected in Naturally Infected Dogs in North America. J. Clin. Microbiol. 43, 5009–5017 (2005).

23. V. Martella, G. Elia, M. Lucente, N. Decaro, E. Lorusso, K. Banyai, M. Blixenkronemoller, N. Lan, R. Yamaguchi, F. Cirone, Genotyping canine distemper virus (CDV) by a hemi-nested multiplex PCR provides a rapid approach for investigation of CDV outbreaks. Vet. Microbiol. 122, 32–42 (2007).

24. Z. Demeter, B. Lakatos, E. A. Palade, T. Kozma, P. Forgách, M. Rusvai, Genetic diversity of Hungarian canine distemper virus strains. Vet. Microbiol. 122, 258–269 (2007).

25. F. Mira, G. Purpari, S. Di Bella, D. Vicari, G. Schirò, P. Di Marco, G. Macaluso, M. Battilani, A. Guercio, Update on canine distemper virus (CDV) strains of Arctic-like lineage detected in dogs in Italy. Vet. Ital. 54, 225–236 (2018).

26. M. Blixenkrone-Möller, V. Svansson, M. Appel, J. Krogsrud, P. Have, C. Orvell, Antigenic relationships between field isolates of morbilliviruses from different carnivores. Arch. Virol. 123, 279–294 (1992).

27. I. K. Visser, V. P. Kumarev, C. Orvell, P. de Vries, H. W. Broeders, M. W. van de Bildt, J. Groen, J. S. Teppema, M. C. Burger, F. G. UytdeHaag, Comparison of two morbilliviruses isolated from seals during outbreaks of distemper in north west Europe and Siberia. Arch. Virol. 111, 149–164 (1990).

28. W. K. Jo, M. Peters, A. Kydyrmanov, M. W. G. Van De Bildt, T. Kuiken, A. Osterhaus, M. Ludlow, The canine morbillivirus strain associated with an epizootic in caspian seals provides new insights into the evolutionary history of this virus. Viruses 11 (2019).

29. M. Marcacci, M. Ancora, I. Mangone, L. Teodori, D. Di Sabatino, F. De Massis, C. Camma’, G. Savini, A. Lorusso, Whole genome sequence analysis of the arctic-lineage strain responsible for distemper in Italian wolves and dogs through a fast and robust next generation sequencing protocol. J. Virol. Methods 202 (2014).

30. D. Di Sabatino, G. Di Francesco, G. Zaccaria, D. Malatesta, L. Brugnola, M. Marcacci, O. Portanti, F. De Massis, G. Savini, L. Teodori, E. Ruggieri, I. Mangone, P. Badagliacca, A. Lorusso, Lethal distemper in badgers (Meles meles) following epidemic in dogs and wolves. Infect. Genet. Evol. 46, 130–137 (2016).

31. M. Tannerfeldt, A. Angerbjorn, Fluctuating resources and the evolution of litter size in the arctic fox. Oikos 83, 545–559 (1998).

32. E. Fuglei, A. Tarroux, Arctic fox dispersal from Svalbard to Canada: One female’s long run across sea ice. Polar Res. 38, 1–7 (2019).

33. R. Gravel, S. Lai, D. Berteaux, Long-term satellite tracking reveals patterns of long-distance dispersal in juvenile and adult Arctic foxes (Vulpes lagopus). R. Soc. Open Sci. 10 (2023).

34. D. T. Shen, J. R. Gorham, Survival of pathogenic distemper virus at 5C and 25C. Vet. Med. Small Anim. Clin. VM SAC 75, 69–72 (1980).

35. E. E. Brandell, E. S. Almberg, P. C. Cross, A. P. Dobson, D. W. Smith, P. J. Hudson, “Infectious Diseases in Yellowstone’s Wolves” in Yellowstone Wolves: Science and Discovery in the World’s First National Park, D. Smith, D. R. Stahler, D. R. MacNulty, Eds. (University of Chicago Press, Chicago, 2020), pp. 121–133.

36. E. S. Almberg, P. C. Cross, D. W. Smith, D. W.. Smith2, Persistence of canine distemper virus in the Greater Yellowstone Ecosystem’s carnivore community. Ecol. Appl. Ecol. Appl. 20, 2058–2074 (2010).

37. P. C. Cross, F. T. van Manen, M. Viana, E. S. Almberg, D. Bachen, E. E. Brandell, M. A. Haroldson, P. J. Hudson, D. R. Stahler, D. W. Smith, F. T. van Manen, M. Viana, E. S. Almberg, D. Bachen, E. E. Brandell, M. A. Haroldson, P. J. Hudson, D. R. Stahler, D. W. Smith, Estimating distemper virus dynamics among wolves and grizzly bears using serology and Bayesian state-space models. Ecol. Evol. 8, 8726–8735 (2018).

38. J. K. Weckworth, B. W. Davis, E. Dubovi, N. Fountain-Jones, C. Packer, S. Cleaveland, M. E. Craft, E. Eblate, M. Schwartz, L. S. Mills, M. Roelke-Parker, Cross-species transmission and evolutionary dynamics of canine distemper virus during a spillover in African lions of Serengeti National Park. Mol. Ecol. 29, 4308–4321 (2020).

39. G. Franzo, L. De Villiers, L. M. Coetzee, M. De Villiers, F. N. Nyathi, M. Garbade, C. Hansen, S. Berjaoui, P. Ripà, A. Lorusso, U. Molini, Unveiling the molecular epidemiology of canine distemper virus in Namibia: An expected pathogen showing an unexpected origin. Heliyon 10, e34805 (2024).

40. A. Wipf, P. Perez-Cutillas, N. Ortega, A. Huertas-López, C. Martínez-Carrasco, M. G. Candela, Geographical Distribution of Carnivore Hosts and Genotypes of Canine Distemper Virus (CDV) Worldwide: A Scoping Review and Spatial Meta-Analysis. Transbound. Emerg. Dis., doi: 10.1155/tbed/6632068 (2025).

41. R. E. Junge, K. Bauman, M. King, M. E. Gompper, A serologic assessment of exposure to viral pathogens and Leptospira in an urban raccoon (Procyon lotor) population inhabiting a large zoological park. J. Zoo Wildl. Med. Off. Publ. Am. Assoc. Zoo Vet. 38, 18–26 (2007).

42. D. W. Smith, D. R. Stahler, M. C. Metz, K. A. Cassidy, E. E. Stahler, E. S. Almberg, M. Rick, YS 24-1 Wolf Restoration in Yellowstone Reintroduction to Recovery – Yellowstone National Park (U.S. National Park Service) (2016). https://www.nps.gov/yell/learn/ys-24-1-wolf-restoration-in-yellowstone-reintroduction-to-recovery.htm.

43. E. S. Almberg, P. C. Cross, A. P. Dobson, D. W. Smith, P. J. Hudson, Parasite invasion following host reintroduction: a case study of Yellowstone’s wolves. Philos. Trans. R. Soc. B Biol. Sci. 367, 2840–2851 (2012).

44. C. H. Gordon, A. C. Banyard, A. Hussein, M. K. Laurenson, J. R. Malcolm, J. Marino, F. Regassa, A. M. E. Stewart, A. R. Fooks, C. Sillero-Zubiri, Canine distemper in endangered Ethiopian wolves. Emerg. Infect. Dis. 21, 824–832 (2015).

45. S. F. Timm, L. Munson, B. A. Summers, K. A. Terio, E. J. Dubovi, C. E. Rupprecht, S. Kapil, D. K. Garcelon, A suspected canine distemper epidemic as the cause of a catastrophic decline in Santa Catalina Island foxes (*Urocyon littoralis catalinae*). J. Wildl. Dis. 45, 333–343 (2009).

46. E. S. Williams, E. T. Thome, M. J. G. Appel, D. W. Belitsky, Canine distemper in black-footed ferrets (*Mustela nigripes*) from Wyoming. J. Wildl. Dis. 24, 385–398 (1988).

47. T. A. Seimon, D. G. Miquelle, T. Y. Chang, A. L. Newton, I. Korotkova, G. Ivanchuk, E. Lyubchenko, A. Tupikov, E. Slabe, D. McAloose, Canine distemper virus: An emerging disease in wild endangered Amur tigers (Panthera tigris altaica). mBio 4, e00410–13 (2013).

48. M. Gilbert, N. Sulikhan, O. Uphyrkina, M. Goncharuk, L. Kerley, E. H. Castro, R. Reeve, T. Seimon, D. McAloose, I. V. Seryodkin, S. V. Naidenko, C. A. Davis, G. S. Wilkie, S. B. Vattipally, W. E. Adamson, C. Hinds, E. C. Thomson, B. J. Willett, M. J. Hosie, N. Logan, M. McDonald, R. J. Ossiboff, E. I. Shevtsova, S. Belyakin, A. A. Yurlova, S. A. Osofsky, D. G. Miquelle, L. Matthews, S. Cleaveland, Distemper, extinction, and vaccination of the amur tiger. Proc. Natl. Acad. Sci. U. S. A. 117, 31954–31962 (2020).

49. N. Ariyama, B. Agüero, B. Bennett, C. Urzúa, F. Berrios, C. Verdugo, V. Neira, Genetic Characterization of Canine morbillivirus (Canine Distemper Virus) Field Strains in Dogs, Chile, 2022–2023. Transbound. Emerg. Dis., doi: 10.1155/2024/9993255 (2024).

50. R. P. Wilkes, Canine Distemper Virus in Endangered Species: Species Jump, Clinical Variations, and Vaccination. Pathogens 12, 57 (2023).

51. E. Anis, A. L. Holford, G. D. Galyon, R. Wilkes, Antigenic analysis of genetic variants of Canine distemper virus. Vet. Microbiol. 219, 154–160 (2018).

52. S. Rendon-Marin, L. F. Higuita-Gutiérrez, J. Ruiz-Saenz, Safety and Immunogenicity of Morbillivirus canis Vaccines for Domestic and Wild Animals: A Scoping Review. Viruses 16, 1078 (2024).

53. F. M. D. Gulland, M. Barbieri, S. Cleaveland, M. Gilbert, A. J. Hall, T. E. Rocke, Vaccination of endangered wildlife as a conservation tool: Hindsights and new horizons in the pandemic era. Biol. Conserv. 300, 110842 (2024).

54. R. K. Maes, A. G. Wise, S. D. Fitzgerald, A. Ramudo, J. Kline, A. Vilnis, C. Benson, A canine distemper outbreak in Alaska: diagnosis and strain characterization using sequence analysis. J. Vet. Diagn. Investig. Off. Publ. Am. Assoc. Vet. Lab. Diagn. Inc 15, 213–220 (2003).

55. Dalerum, Fredrik, Shults, Brad, Kunkel, Kyran, A Serologic Survey for Antibodies to Three Canine Viruses in Wolverines (*Gulo gulo*) from the Brooks Range, Alaska. J. Wildl. Dis. 41 (2005).

56. J. A. Nasir, R. A. Kozak, P. Aftanas, A. R. Raphenya, K. M. Smith, F. Maguire, H. Maan, M. Alruwaili, A. Banerjee, H. Mbareche, B. P. Alcock, N. C. Knox, K. Mossman, B. Wang, J. A. Hiscox, A. G. McArthur, S. Mubareka, A comparison of whole genome sequencing of sars-cov-2 using amplicon-based sequencing, random hexamers, and bait capture. Viruses 12, 895 (2020).

57. T. S. Korneliussen, A. Albrechtsen, R. Nielsen, ANGSD: Analysis of Next Generation Sequencing Data. BMC Bioinformatics 15, 1–13 (2014).

58. A. L. Frisk, M. König, A. Moritz, W. Baumgärtner, Detection of canine distemper virus nucleoprotein RNA by reverse transcription-PCR using serum, whole blood, and cerebrospinal fluid from dogs with distemper. J. Clin. Microbiol. 37, 3634–3643 (1999).

59. J. Quick, N. D. Grubaugh, S. T. Pullan, I. M. Claro, A. D. Smith, K. Gangavarapu, G. Oliveira, R. Robles-Sikisaka, T. F. Rogers, N. A. Beutler, D. R. Burton, L. L. Lewis-Ximenez, J. Goes de Jesus, M. Giovanetti, S. Hill, A. Black, T. Bedford, M. W. Carroll, M. Nunes, L. C. Alcantara, E. C. Sabino, S. A. Baylis, N. Faria, M. Loose, J. T. Simpson, O. G. Pybus, K. G. Andersen, N. J. Loman, Multiplex PCR method for MinION and Illumina sequencing of Zika and other virus genomes directly from clinical samples. bioRxiv 12, 098913 (2017).

60. D. M. Gohl, J. Garbe, P. Grady, J. Daniel, R. H. B. Watson, B. Auch, A. Nelson, S. Yohe, K. B. Beckman, A rapid, cost-effective tailed amplicon method for sequencing SARS-CoV-2. BMC Genomics 21, 1–10 (2020).

61. H. Li, Aligning sequence reads, clone sequences and assembly contigs with BWA-MEM. ArXiv Prepr. 1303.3997, 1–3 (2013).

62. H. Li, B. Handsaker, A. Wysoker, T. Fennell, J. Ruan, N. Homer, G. Marth, G. Abecasis, R. Durbin, The Sequence Alignment/Map format and SAMtools. Bioinformatics 25, 2078–2079 (2009).

63. N. D. Grubaugh, K. Gangavarapu, J. Quick, N. L. Matteson, J. G. De Jesus, B. J. Main, A. L. Tan, L. M. Paul, D. E. Brackney, S. Grewal, N. Gurfield, K. K. A. Van Rompay, S. Isern, S. F. Michael, L. L. Coffey, N. J. Loman, K. G. Andersen, An amplicon-based sequencing framework for accurately measuring intrahost virus diversity using PrimalSeq and iVar. Genome Biol. 20, 1–19 (2019).

64. A. R. Quinlan, I. M. Hall, BEDTools: A flexible suite of utilities for comparing genomic features. Bioinformatics 26, 841–842 (2010).

65. D. Posada, jModelTest: Phylogenetic model averaging. Mol. Biol. Evol. 25, 1253–1256 (2008).

66. S. Tavaré, Some probabilistic and statistical problems in the analysis of DNA sequences. Am. Math. Soc. Lect. Math. Life Sci. 17, 57–86 (1986).

67. A. Rambaut, T. T. Lam, L. M. Carvalho, O. G. Pybus, Exploring the temporal structure of heterochronous sequences using TempEst (formerly Path-O-Gen). Virus Evol. 2, vew007 (2016).

68. A. J. Drummond, M. A. Suchard, D. Xie, A. Rambaut, Bayesian phylogenetics with BEAUti and the BEAST 1.7. Mol. Biol. Evol. 29, 1969–1973 (2012).

69. A. J. Drummond, A. Rambaut, B. Shapiro, O. G. Pybus, Bayesian coalescent inference of past population dynamics from molecular sequences. Mol. Biol. Evol. 22, 1185–1192 (2005).

70. A. J. Drummond, G. K. Nicholls, A. G. Rodrigo, W. Solomon, Estimating mutation parameters, population history and genealogy simultaneously from temporally spaced sequence data. Genetics 161, 1307–1320 (2002).

71. A. J. Drummond, S. Y. W. Ho, M. J. Phillips, A. Rambaut, Relaxed phylogenetics and dating with confidence. PLoS Biol. 4, 699–710 (2006).

72. J. R. Gog, B. T. Grenfell, Dynamics and selection of many-strain pathogens. Proc. Natl. Acad. Sci. U. S. A. 99, 17209–17214 (2002).

73. E. M. Volz, K. Koelle, T. Bedford, Viral Phylodynamics. PLoS Comput. Biol. 9, e1002947 (2013).

74. M. D. Karcher, L. M. Carvalho, M. A. Suchard, G. Dudas, V. N. Minin, Estimating effective population size changes from preferentially sampled genetic sequences. PLoS Comput. Biol. 16, e1007774 (2020).

75. M. Viana, J. A. Benavides, A. Broos, D. I. Loayza, R. Niño, J. Bone, A. da Silva Filipe, R. Orton, W. V. Bazan, J. Matthiopoulos, D. G. Streicker, Effects of culling vampire bats on the spatial spread and spillover of rabies virus. Sci. Adv. 9 (2023).

76. R. C. Griffiths, S. Tavaré, Sampling theory for neutral alleles in a varying environment. Philos. Trans. R. Soc. Lond. B. Biol. Sci. 344, 403–410 (1994).

77. J. Wallinga, M. Lipsitch, How generation intervals shape the relationship between growth rates and reproductive numbers. Proc. R. Soc. B Biol. Sci. 274, 599–604 (2007).

78. N. C. Grassly, C. Fraser, Mathematical models of infectious disease transmission. Nat. Rev. Microbiol. 6, 477–487 (2008).

79. N. M. Ferguson, D. A. T. Cummings, S. Cauchemez, C. Fraser, S. Riley, A. Meeyai, S. Iamsirithaworn, D. S. Burke, Strategies for containing an emerging influenza pandemic in Southeast Asia. Nature 437, 209–214 (2005).

80. C. Fraser, C. A. Donnelly, S. Cauchemez, W. P. Hanage, M. D. Van Kerkhove, T. D. Hollingsworth, J. Griffin, R. F. Baggaley, H. E. Jenkins, E. J. Lyons, T. Jombart, W. R. Hinsley, N. C. Grassly, F. Balloux, A. C. Ghani, N. M. Ferguson, A. Rambaut, O. G. Pybus, H. Lopez-Gatell, C. M. Alpuche-Aranda, I. B. Chapela, E. P. Zavala, D. Ma. Espejo Guevara, F. Checchi, E. Garcia, S. Hugonnet, C. Roth, Pandemic potential of a strain of influenza A (H1N1): Early findings. Science 324, 1557–1561 (2009).

81. H. L. Del Puerto, A. C. Vasconcelos, L. Moro, F. Alves, G. F. Braz, A. S. Martins, Canine distemper virus detection in asymptomatic and non vaccinated dogs. Pesqui. Veterinária Bras. 30, 139–144 (2010).

82. S. Newbury, L. Larson, R. Schultz, “Canine Distemper Virus” in Infectious Disease Management in Animal Shelters, L. Miller, K. Hurley, Eds. (Wiley-Blackwell, 2009).

83. J. Graham, R. P. Marini, G. J. Fox, “Ferrets: Distemper” in Clinical Veterinary Advisor: Birds and Exotic Pets, J. Mayer, T. M. Donnelly, Eds. (Elsevier Inc., 2012), pp. 444–445.

84. A. J. Drummond, A. Rambaut, BEAST: Bayesian evolutionary analysis by sampling trees. BMC Evol. Biol. 7, 1–8 (2007).

85. H. Wickham, Ggplot2: Elegant Graphics for Data Analysis (Springer-Verlag New York, New York, 2016; http://link.springer.com/10.1007/978-0-387-98141-3).

