## Supplement for "Persistent, Multi-Species Outbreaks and Long-Range Transmissions of a Measles-Like Virus"

### 1 Supplement

| Species | Year | No. of samples processed | CDV detected | Sequences >70% coverage | Sequences 45–70% coverage | Sequences 30–45% coverage | Sequences 1–30% coverage | GMU numbers |
|---|---|---|---|---|---|---|---|---|
| Arctic fox | 2012 | 17 | 9 | 4 | 3 | 2 | 0 | 26 |
| Arctic fox | 2013 | 11 | 4 | 3 | 0 | 1 | 0 | 26 |
| Arctic fox | 2014 | 21 | 11 | 3 | 2 | 3 | 0 | 26 |
| Arctic fox | 2016 | 5 | 1 | 0 | 0 | 0 | 1 | 26 |
| Arctic fox | 2017 | 3 | 0 | 0 | 0 | 0 | 0 | 26 |
| Arctic fox | 2021 | 45 | 29 | 9 | 8 | 9 | 3 | 22 |
| Red fox | 2012 | 4 | 0 | 0 | 0 | 0 | 0 | 14, 20, 25, 26 |
| Red fox | 2014 | 2 | 0 | 0 | 0 | 0 | 0 | 8, 20, 26 |
| Red fox | 2015 | 6 | 0 | 0 | 0 | 0 | 0 | 14, 26 |
| Red fox | 2016 | 17 | 0 | 0 | 0 | 0 | 0 | 14, 17, 18, 20, 26 |
| Red fox | 2017 | 8 | 0 | 0 | 0 | 0 | 0 | 14, 20, 23, 26 |
| Red fox | 2021 | 27 | 4 | 0 | 0 | 0 | 1 | 20,26 |
| Domestic dog | 2021 | 2 | 2 | 0 | 1 | 1 | 0 | 22 |
| Gray wolf | 2013 | 1 | 0 | 0 | 0 | 0 | 0 | AK |
| Gray wolf | 2015 | 23 | 0 | 0 | 0 | 0 | 0 | 1, 2, 13, 20, 25, 26 |
| Gray wolf | 2017 | 1 | 1 | 0 | 1 | 0 | 0 | YNP |
| Black bear | 2016 | 2 | 2 | 0 | 0 | 2 | 0 | 14 |
| Black bear | 2017 | 7 | 7 | 0 | 0 | 3 | 4 | 1,13, 14, 20 |
| Brown bear | 2015 | 2 | 2 | 0 | 0 | 1 | 1 | 9,26 |
| Brown bear | 2016 | 4 | 4 | 0 | 0 | 1 | 3 | 20 |
| Brown bear | 2017 | 4 | 2 | 0 | 0 | 1 | 1 | 26 |
| Wolverine | 2015 | 2 | 1 | 0 | 0 | 0 | 1 | 20 |
| Wolverine | 2016 | 12 | 5 | 0 | 0 | 1 | 3 | 12, 18, 20, 22, 26 |
| Marten | 2015 | 1 | 1 | 0 | 0 | 0 | 0 | 18 |
| Marten | 2016 | 1 | 0 | 0 | 0 | 0 | 0 | 20 |
| River otter | 2015 | 1 | 0 | 0 | 0 | 0 | 0 | 5 |
| River otter | 2016 | 2 | 1 | 0 | 0 | 0 | 1 | 15,19 |
| Lynx | 2015 | 1 | 0 | 0 | 0 | 0 | 0 | 15 |

| | | | | | | | | |
|---|---|---|---|---|---|---|---|---|
| Lynx | 2016 | 1 | 0 | 0 | 0 | 0 | 0 | 15 |
| Coyote | 2015 | 2 | 0 | 0 | 0 | 0 | 0 | 18,20 |
| **Total** | | **235** | **86** | **19** | **15** | **25** | **19** | |

**Table S1. The number of total samples, CDV-positive samples, and sequenced CDV genomes (with high, and mid- to low coverage) obtained in this study per species and sample year.** Here we show all processed samples per species and sample year from a large CDV screening effort in Alaskan carnivores and a Yellowstone gray wolf. Game management units (GMUs), Yellowstone National Park (YNP), canine distemper virus (CDV), Alaska (AK). The species' common and latin names were: Arctic fox (*Vulpes lagopus*), red fox (*Vulpes vulpes*), domestic dog (*Canis lupus familiaris*), gray wolf (*Canis lupus*), black bear (*Ursus americanus*), brown bear (*Ursus arctos*), wolverine (*Gulo gulo*), marten (*Martes americana*), river otter (*Lontra canadensis*), lynx (*Lynx canadensis*), and coyote (*Canis latrans*).

| Sample ID | No. of reads | % mapping to CDV reference (no. of reads) | Coverage depth of CDV genome | % of CDV reference covered | % reads mapped to Arctic fox reference genome |
|---|---|---|---|---|---|
| 255 | 3,945,034 | 0 (2) | 0.019 | 1.87 | 99.73 |
| 275 | 3,968,150 | 0.1 (4050) | 36.86 | 94.34 | 99.68 |
| 276 | 8,883,734 | 0 (15) | 0.14 | 1.93 | 99.79 |
| 330 | 8,876,016 | 0 (262) | 2.43 | 21.05 | 99.73 |
| 345 | 4,946,936 | 0 (18) | 0.17 | 1.89 | 99.64 |
| 404 | 17,926,436 | 0 (4) | 0.04 | 3.85 | 99.82 |
| 419 | 5,541,316 | 0.01 (449) | 3.68 | 29.55 | 99.75 |
| 441 | 4,053,078 | 0 (2) | 0.02 | 1.93 | 99.66 |

**Table S2. Sequencing results using myBaits.**

**Supplementary File S1. PCR1 primer sequences with 4 pools indicated.** The base pair start and stop point of each amplicon is indicated in relation to the reference genomes used for primer design (sample 275 filled in with KF914669). The first part of each primer sequence (TCGTCGGCAGCGTCAGATGTGTATAAGAGACAG for left primers and GTCTCGTGGGCTCGGAGATGTGTATAAGAGACAG for right primers) are adapter sequences needed for the second PCR. The second part of each primer is the CDV-specific primer sequence generated by Primal Scheme.

**Supplementary File S2. PCR2 barcoding primer sequences.** All forward i7 primers and reverse i5 primers used for unique dual indexing are indicated. The first part of all primers (CAAGCAGAAGACGGCATACGAGAT for i7 primers and AATGATACGGCGACCACCGAGATCTACAC for i5 primers) are Illumina flow-cell adapters. The 10 base pairs following the Illumina flow-cell adapters are the unique index sequence. The last part of each primer (GTCTCGTGGGCTCGG for i7 primers and TCGTCGGCAGCGTC for i5 primers) are adapter sequences that are complementary to those in the PCR1 primers.

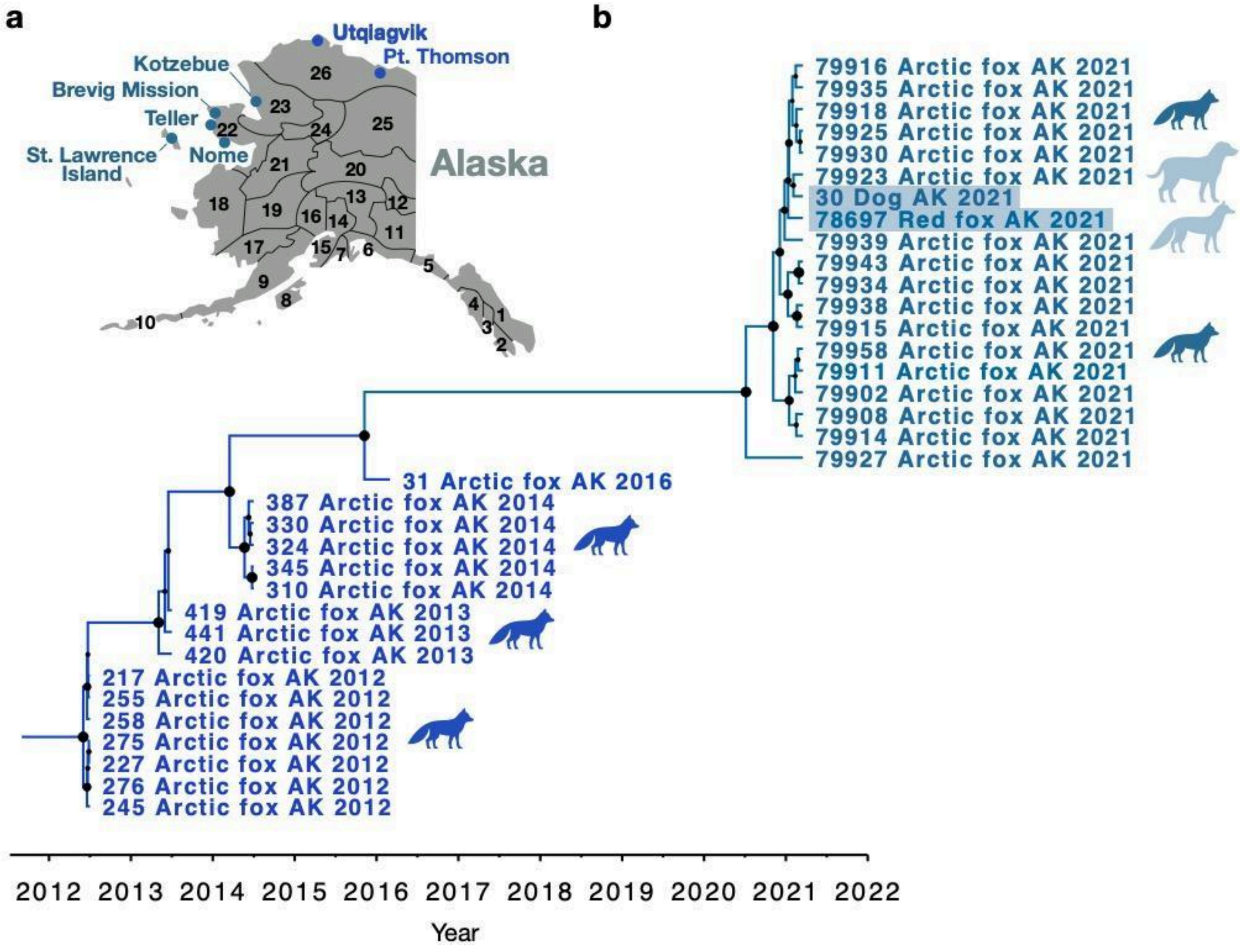


**Figure S1. Additional support for canine distemper virus (CDV) persistence in Arctic foxes, and possible cross-species transmission of CDV between Arctic fox and red fox during the 2021 outbreak.** This figure is almost the same as Figure 1 in the main text except with the addition of partial genomes with low coverage from an Arctic fox in 2016 and a red fox in 2021. (a) Map of Alaska with game management unit (GMU) locations, where most of the sampling has been done in Alaska. The colors of the samples in (b) match the colors on the map and indicate where the samples have been collected. (b) A dated phylogenetic (maximum clade credibility) tree of 33 CDV full or partial genomes from Arctic foxes, one domestic dog (highlighted in light blue), and one red fox (highlighted in light blue) sampled in 2012–2014 (blue) and 2021 (steel blue) in Alaska. The horizontal positioning of each node (along the timeline) represents the estimated median of the time of divergence in years (from the most recent common ancestor [MRCA]). The size of the black dots on the nodes represents the size of the posterior probability (PP) with the maximum size (PP = 1) equivalent to the size of the black dot at the root of the tree (January, 2012). All genomes had a coverage of 45% or higher, except the sequence from the red fox (4.4% coverage) and an Arctic fox sampled in 2016 (20% coverage). Despite its lower coverage, the viral sequence from the red fox clusters within the 2021 clade, lending support of possible cross-species transmission between red fox and Arctic fox. The 2016 Arctic fox sample provides further support for possible long-term persistence of CDV in Arctic foxes.

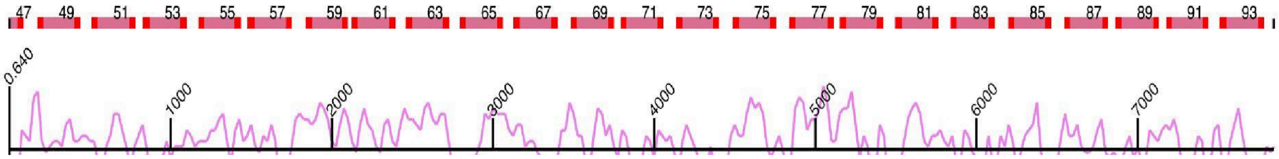


**Figure S2. A schematic showing how the multiplex PCR primers sit along the length of the CDV genome.** The number of each amplicon is indicated. The bright red sections indicate the primers and the lighter red segments indicate the amplicons. All of the odd numbered primers (along the top) go into one pool and all of the even numbered primers (along the bottom) go into a second pool, so that neighboring amplicons do not overlap within the same pool.

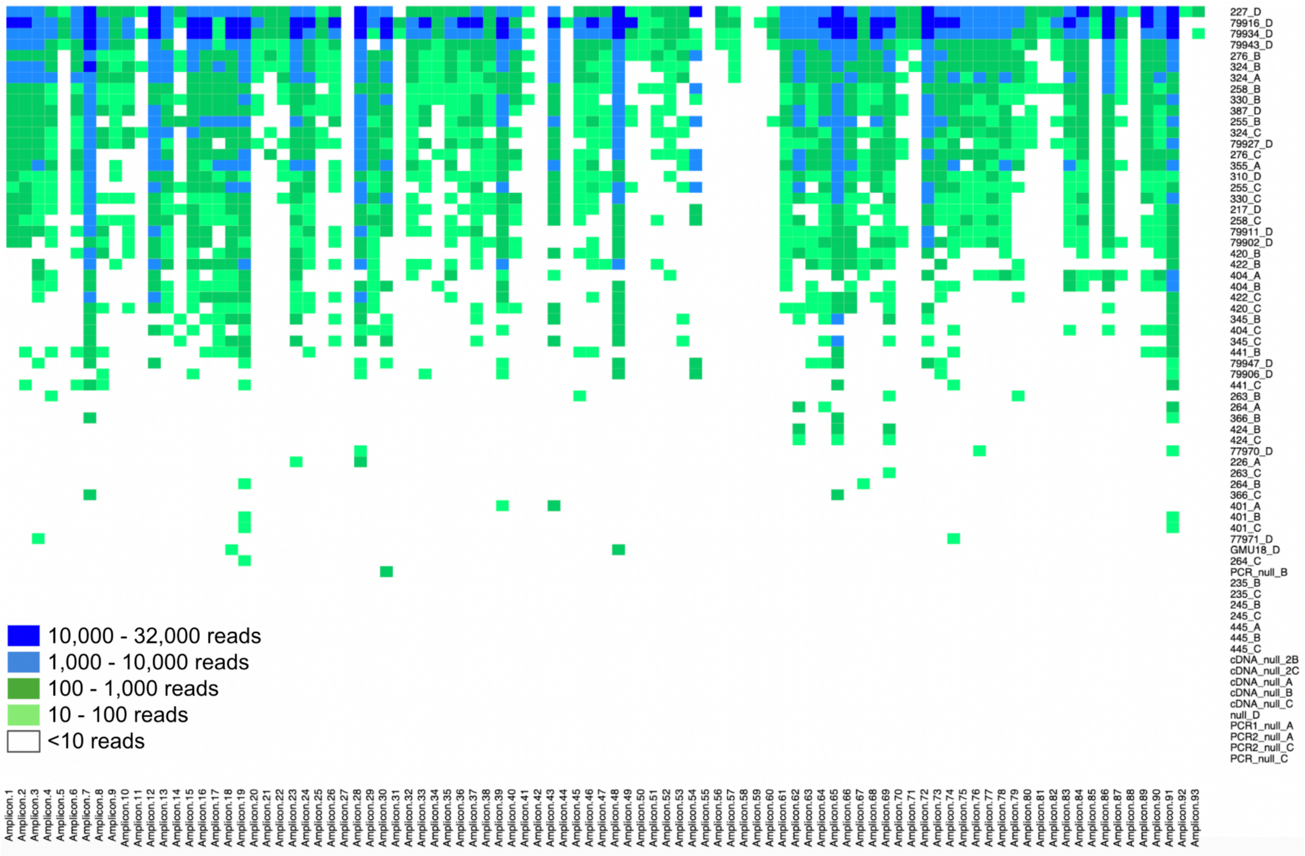


**Figure S3. A heat map indicating the number of reads generated for each amplicon for each sample (2 primer pools used).**

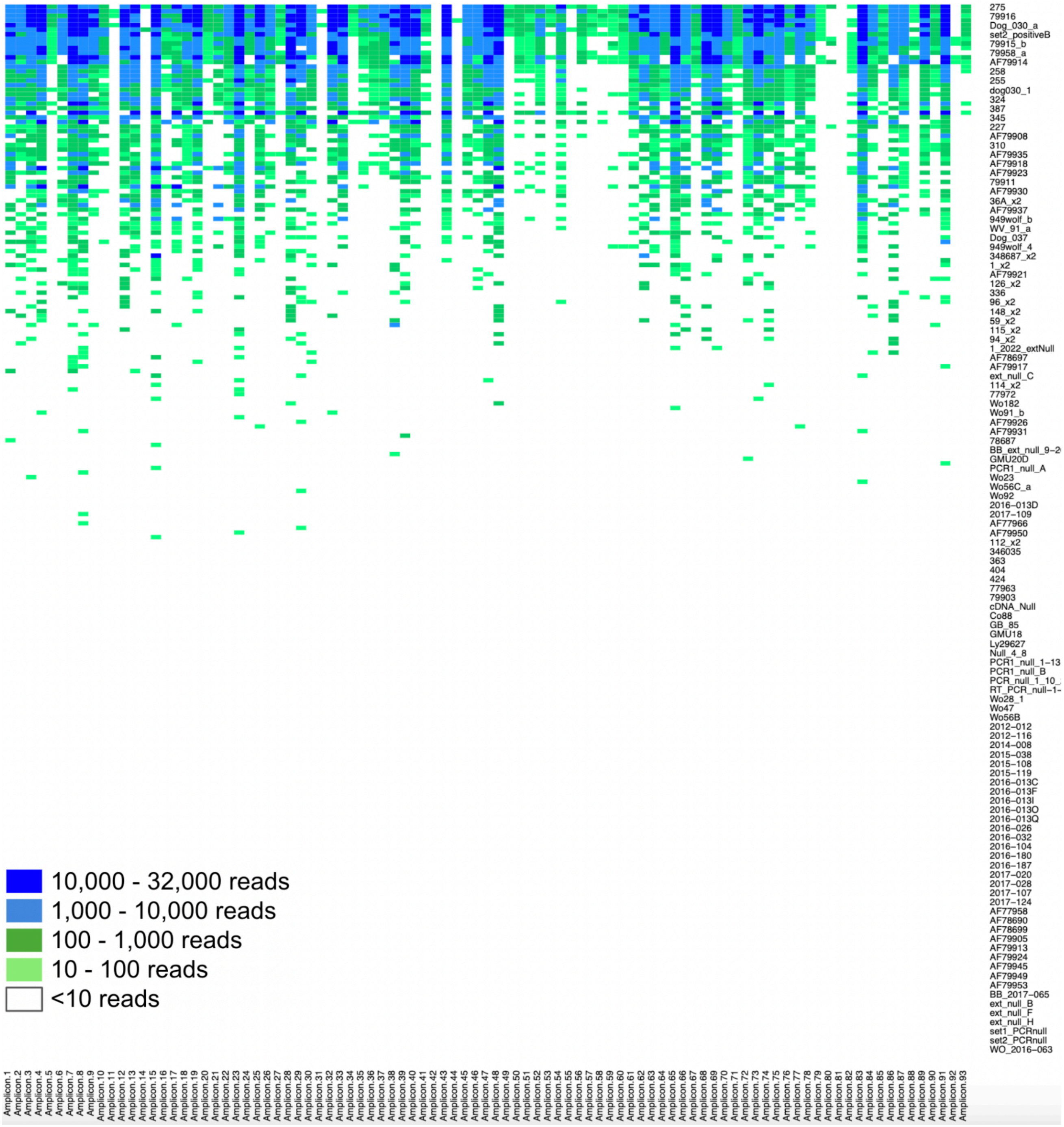


**Figure S4. A heat map indicating the number of reads generated for each amplicon for each sample (4 primer pools used).** Although 227 samples were processed, only the top 144 are shown here, as the remaining samples showed less than 10 reads.

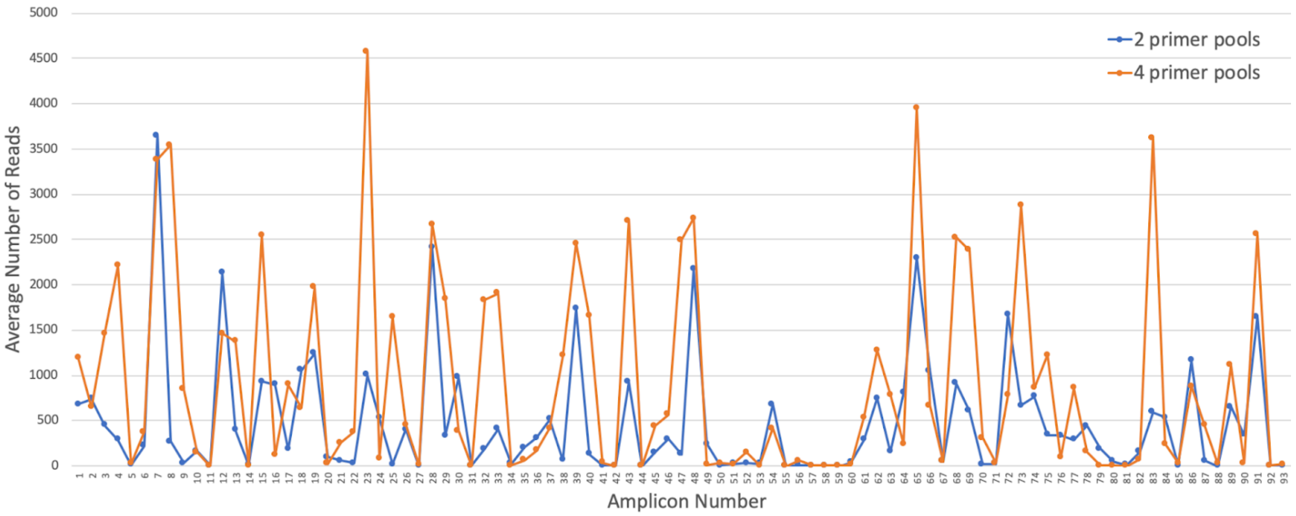


**Figure S5. Average number of reads per sample for each of the 93 amplicons.**
